# Optimization of an adeno-associated viral vector for epidermal keratinocytes *in vitro* and *in vivo*

**DOI:** 10.1101/2024.04.15.589645

**Authors:** Qi Shen, Shogo Suga, Yuta Moriwaki, Zening Du, Emi Aizawa, Mutsumi Okazaki, Juan Carlos Izpisua Belmonte, Yusuke Hirabayashi, Keiichiro Suzuki, Masakazu Kurita

## Abstract

**Background:** Local gene therapies, including *in vivo* genome editing, are highly anticipated for the treatment of genetic diseases in skin, especially the epidermis. While the adeno-associated virus (AAV) is a potent vector for *in vivo* gene delivery, the lack of efficient gene delivery methods has limited its clinical applications.

**Objective:** To optimize the AAV gene delivery system with higher gene delivery efficiency and specificity for epidermis and keratinocytes (KCs), using AAV capsid and promoter engineering technologies.

**Methods:** AAV variants with mutations in residues reported to be critical to determine the tropism of AAV2 for KCs were generated by site-directed mutagenesis of AAVDJ. The infection efficiency and specificity for KCs of these variants were compared with those of previously reported AAVs considered to be suitable for gene delivery to KCs *in vitro* and *in vivo*. Additionally, we generated an epidermis-specific promoter using the most recent short-core promoter and compared its specificity with existing promoters.

**Results:** A novel AAVDJ variant capsid termed AAVDJK2 was superior to the existing AAVs in terms of gene transduction efficiency and specificity for epidermis and KCs *in vitro* and *in vivo*. A novel tissue-specific promoter, termed the K14 SCP3 promoter, was superior to the existing promoters in terms of gene transduction efficiency and specificity for KCs.

**Conclusion:** The combination of the AAVDJK2 capsid and K14 SCP3 promoter improves gene delivery to epidermis *in vivo* and KCs *in vitro*. The novel AAV system may benefit experimental research and development of new epidermis-targeted gene therapies.

## Introduction

Epidermis, composed dominantly of keratinocytes (KCs), plays an important role in skin homeostasis, wound healing, and tissue regeneration [1][2]. Many skin disorders are caused by genetic dysfunctions and/or mutations [3]. Despite the easy accessibility of the skin and the urgent medical needs, the application of gene therapy to skin-related diseases has received little attention because of the low achievable efficiency of gene transduction [4].

Adeno-associated viruses (AAVs) are useful vectors for *in vivo* gene transduction and are clinically used as replacement therapies to treat diseases such as lipoprotein lipase deficiency, spinal muscular atrophy, retinal dystrophy, and hemophilia [5]. Recently, they have become promising vectors for the development of potent therapeutics targeting localized morbidities [2][6][7] and therapeutic *in vivo* genome editing [8].

Each serotype (*i.e.*, capsid structure variant) of AAV has a specific cell/tissue tropism, and engineering of the capsid, the protein shell that surrounds the viral genome, allows for increased gene transduction efficiency and specificity for a particular cell/tissue, thereby minimizing unspecific off-target effects [9].

The use of AAVs for KCs has mostly been described in the context of *in vitro* applications. Sallach *et al.* generated a highly efficient engineered AAV2 through tropism engineering by directed evolution of AAV2 [10] and identified a variant targeting αvβ8 integrin on KCs. Furthermore, Melo *et al.* reported that AAVDJ, a hybrid serotype, outperforms previously reported AAV2 and AAV6 in terms of transduction efficiency of KCs *in vitro* [11].

We reasoned that modification of AAVDJ with an αvβ8 integrin-targeting variant would generate an optimized hybrid AAV vector for *in vivo* gene transduction of the epidermis. In the current study, the relative superiority of the novel hybrid AAV over existing AAVs considered suitable for KCs was evaluated *in vitro* and *in vivo*. The performance of tissue-specific promoters, including originals, was also evaluated in terms of gene transduction efficiency and specificity *in vitro* to optimize the novel AAV system useful for the development of epidermis-targeted gene therapies.

## Materials and methods

### Approval for animal studies and biosafety

All animal experiments were approved by the Animal Research Committee of the University of Tokyo (IRB approval number Med-P21-024). All genetic modification experiments were approved by the Biosafety Committee of the University of Tokyo (IBC approval number 37-5).

### Generation of AAV capsid plasmids by mutagenesis

Site-directed mutagenesis was performed with a QuikChange Lightning Site-Directed Mutagenesis Kit (Agilent technologies, Inc., Santa Clara, CA) to generate AAV2 and AAVDJ using keratinocyte-targeting variants [10][12] in pXR2 (encoding AAV2 Rep-Cap) and pAAVDJ plasmids (Cell Biolabs, Inc., San Diego, CA), respectively.

### AAV plasmid construction

The pAAV-CAG-GFPNLS plasmid was prepared by modification of pAAV-CAG-GFP (Plasmid #37825) from Addgene using an In-Fusion® HD Cloning Kit (Clontech, Mountain View, CA). The human *KRT14* enhancer and promoter sequence [13] [14] was amplified using human genomic DNA from HEK293 cells with PrimeSTAR GXL DNA polymerase (Takara Bio Inc., Shiga, Japan). The human *KRT16* pseudogene 5 (K16P5) enhancer and promoter sequence was amplified from pRZ-hKeratin (Cat# SR900CS-1, System Biosciences). The SCP3 core promoter [15] was synthesized by Genewiz. To generate the pAAV-K14-GFPNLS plasmid, 2 kb of the *KRT14* (K14) enhancer/promoter replaced the CAG promoter of pAAV-CAG-GFPNLS using the In-Fusion HD Cloning kit. To generate the pAAV-K14short-GFPNLS plasmid, 0.7 kb of the *KRT14* enhancer and 0.3 kb of the *KRT14* minimal promoter [14] replaced the CAG promoter of pAAV-CAG-GFPNLS using the In-Fusion HD Cloning kit. To generate the pAAV-K14SCP3-GFPNLS plasmid, 0.7 kb of the *KRT14* enhancer and 0.1 kb of the SCP3 core promoter replaced the CAG promoter of pAAV-CAG-GFPNLS using the In-Fusion HD Cloning kit. To generate the pAAV-K16P5-GFPNLS plasmid, 2 kb of the K16P5 enhancer-promoter replaced the CAG promoter of pAAV-CAG-GFPNLS using the In-Fusion HD Cloning kit. To generate the pAAV-K16P5short-GFPNLS plasmid, 0.7 kb of the K16P5 enhancer and 0.3 kb of the K16P5 minimal promoter replaced the CAG promoter of pAAV-CAG-GFPNLS using the In-Fusion HD Cloning kit. To generate the pAAV-K16P5SCP3-GFPNLS plasmid, 0.7 kb of the K16P5 enhancer and 0.1 kb of the SCP3 core promoter replaced the CAG promoter of pAAV-CAG-GFPNLS using the In-Fusion HD Cloning kit.

### AAV production

AAVs were prepared using 293AAV cells (Cell Biolabs, Inc.) as described previously [2] [16] with minor modifications (calcium phosphate transfection followed by CsCl gradient purification). The virus titer was determined by qPCR using primers ITR-F, 5ʹ-GGAACCCCTAGTGATGGAGTT-3ʹ and ITR-R, 5ʹ-CGGCCTCAGTGAGCGA-3ʹ.

### Mouse keratinocyte isolation and culture

Mouse keratinocytes were isolated from the back skin of 3–5-week-old female C57BL/6J Jcl mice which were purchased from Nippon Bio-Supp (Japan). The skin specimens were washed three times in phosphate-buffered saline (PBS), finely shredded with scissors, and incubated with 0.25% trypsin and 0.02% ethylenediaminetetraacetic acid (EDTA) in PBS for 16–24 h at 4 °C. Trypsinized skin pieces were suspended in F medium (3:1 [v/v] Ham’s F12 nutrient mixture-DMEM, high glucose [Wako Pure Chemical Industries, Ltd., Osaka, Japan]) supplemented with 5% FBS, 0.4 μg/ml hydrocortisone (Sigma-Aldrich, St Louis, MO), 5 μg/ml insulin (Sigma), 8.4 ng/ml cholera toxin (Wako), 10 ng/ml EGF (Sigma), 24 μg/ml adenine (Sigma), 100 U/ml penicillin, 100 μg/ml streptomycin (Wako), and 10 μM Rho-kinase inhibitor Y27632 (Selleck Chemicals, Houston, TX) [2]. The epidermis is peeled off, and the remaining epidermal cells adhering to the superficial side of the dermis are further peeled off with square tweezers to recover the epidermal cells. The cells are isolated by centrifugation and maintained on mitomycin C-treated 3T2-J2 feeder cells in F medium.

### Human keratinocyte culture

Normal human epidermal keratinocytes (adult donor, pooled, Product code: C12006; PromoCell, Heidelberg, Germany) were maintained on 3T3-J2 feeder cells under the same conditions used for mouse keratinocytes.

### Mouse adipose-derived stromal cell isolation culture

Mouse adipose-derived stromal cells were isolated from subcutaneous groin-lumber fat pads of 3–5-week-old female C57BL/6JJcl mice which were purchased from Nippon Bio-Supp (Japan). Briefly, adipose tissue was enzymatically digested as described previously [17]. The stromal vascular fraction was isolated by centrifugation and maintained in complete growth medium consisting of DMEM containing 4.5 g/L glucose, 110 mg/L sodium pyruvate, and 4 mM L-glutamine supplemented with 10% (v/v) heat-inactivated fetal bovine serum, 1:100 (v/v) MEM non-essential amino acid solution (Wako), and 1:100 (v/v) GlutaMAX supplement (Wako) [2].

### Human dermal fibroblast culture

Normal human dermal fibroblasts (juvenile foreskin, Product Code: C12300; PromoCell, Heidelberg, Germany) were maintained under the same conditions used for mouse adipose-derived stromal cells.

### Analysis of gene transduction efficiency of keratinocytes

Keratinocytes were cultured on feeders in 24-well plates. At 50% confluence, the medium was changed to medium containing 1×10^9^ GC/well of AAVs for four technical replicates (day 0). On the day of assessment, phase contrast and fluorescence images were obtained at four positions. On the final day of each assessment, cells were fixed with 4% paraformaldehyde (PFA), stained with 4ʹ,6-diamidino-2-phenylindole (DAPI), and imaged. The efficiency of gene transduction was calculated as the GFPNLS-positive cell number per area of keratinocyte colonies defined in phase contrast images. ImageJ software was used to quantify the areas.

### Analysis of gene transduction efficiency of mesenchymal cells

Mesenchymal cells were seeded in 24-well plates. At 50% confluence, the medium was changed to medium containing 1×10^9^ GC/well of AAVs for four technical replicates (day 0). On the day of assessment, phase contrast and fluorescent images were obtained at four positions. On the final day of each assessment, cells were fixed with 4% PFA, stained with DAPI, and images. The number of positive cells per fluorescence image (area: 1.99 mm^2^) was counted.

### Animals

For *in vivo* experiments, 4-week-old female C57BL/6 mice were purchased from Nippon Bio-Supp (Japan) and used between the ages of 4 and 6 weeks.

### Evaluation of the nanoparticle distribution after intradermal injection

Original silica nanoparticles (1 mg/ml, Sicastar^®^-green F, 30 nm diameter; Micromod Partikeltechnologies, Germany) were diluted 20-fold in PBS. Under inhalation anesthesia, 20 µl of the solution was injected intradermally using a 1 ml syringe with a 29 G needle (SS-10M2913; Terumo Corp., Tokyo, Japan). To ensure intradermal injection into the thin mouse dermis, all injections were performed under a high-resolution surgical microscope (MM100-YOH; Mitaka Kohki, Tokyo, Japan). At 0, 1, and 3 hours after injection, a portion of the skin was collected and immediately embedded in OCT compound. Samples were sectioned using a cryofilm (Cryofilm type 2C(9), 2.5 cm C-FP094; Section-Lab, Hiroshima, Japan) with a previously described method [18] at 200 µm intervals. Nuclei were counterstained with DAPI. Samples were imaged with a slide scanner (VS200-FL; Olympus, Tokyo, Japan).

### Analysis of gene transduction efficiency *in vivo*

Mixtures of 1×10^11^ GC (genome copies) of AAVDJK2-CAG-GFPNLS and 1×10^11^ GC AAVDJ-CAG-mCherryNLS were prepared as a 35 µl solution and injected intradermally into mouse back skin under an operative microscope (n=6). Two days later, the regional skin was collected with the assistance of a stereoscope. The samples were fixed in 4% PFA for 24 h, immersed in 30% sucrose for 2 days, and then embedded in OCT compound. Samples were sectioned with the cryofilm at 200 µm intervals. The center of gene transduction was identified by observation under a fluorescence microscope. The sections were further processed for nuclear staining and imaging.

For machine-assisted quantitative analyses, an 800×800 µm rectangular frame was set at the central site of gene transfer. With the guidance of nearby HE-stained sections, the border between each skin layer (epidermis, dermis, superficial fat, cutaneous muscle, and deep fat) was defined manually.

For the comparative analysis of the *in vivo* gene transduction efficiency of AAVDJK2-CAG-GFPNLS and AAVDJK2-K14 SCP3-GFPNLS, 35 µl of 1×10^11^ GC solution was tested in 3 animals each using the same technique. Cryosections were processed for immunohistochemistry using cytokeratin 14 antibody (ab181595, 1:500; Abcam) as previously described [2].

### Automated analysis of gene transduction efficiency

To unbiasedly extract nuclei from DAPI-stained images of each skin layer (epidermis, dermis, superficial fat, cutaneous muscle, and deep fat), DAPI-stained images were binarized by multiple thresholding steps. Initially, the images were binarized by the threshold set to 0.1875 × the maximum intensity of the DAPI-stained images (*MIPdapi*) of each skin layer. Individual objects in the binarized images were labeled and objects within the size range of [10, 100] pixels were defined as nuclei. For the second cycle, we masked the areas defined as nuclei in the first cycle, and the same procedure was performed with the threshold set to 0.2 × *MIPdapi*. Then, newly defined objects were merged with the nuclear regions defined in the first cycle. By repeating the process 15 times while incrementally adding 0.0125 to the multiplier for *MIPdapi*, the nuclear regions were segmented. Next, to determine whether the nuclei defined above were positive for mCherry or GFP, we set a threshold for binarization of each image so that the area of the binarized image was 4% of the total, and binarized images accordingly. Nuclei with more than five mCherry-positive pixels were defined as nuclei from AAVDJ-CAG-mCherryNLS-infected cells, and nuclei with more than five GFP-positive pixels were defined as nuclei from AAVDJK2-CAG-GFPNLS-infected cells. We visually verified that the binarization defined GFP-and mCherry-positive cells correctly. The numbers of infected cells were counted in each region and layer. For each sample, 30 sections from six mice were analyzed.

### Flow cytometric analysis

Primary mouse KCs and primary mASCs were seeded in 24-well plates. At 50% confluence, the medium was replaced with medium containing 1×10^9^ GC of AAVs per well for four biological replicates. The medium was changed on days 1 and 3. On day 4, single cell suspensions of KCs and ASCs were prepared. Each suspension was prepared by passing the cells through a 35 µm filter using Dulbecco’s PBS containing 5% fetal bovine serum (FBS) and 2 mM EDTA. For cell analysis, a FACSMelody Cell Sorter (BD Biosciences, San Jose, CA) and FlowJo (BD) were used. Single cells were gated sequentially using two-parameter FSC-A versus SSC-A, SSC-H versus SSC-W, and FSC-H versus FSC-W dot plots, and then a one-parameter histogram of GFP fluorescence intensities was plotted for the selected population.

### AAV production analysis

To determine the influence of the AAV type on production, AAV2-CAG-GFPNLS, AAVDJ-CAG-GFPNLS, AAVDJK2-CAG-GFPNLS, AAVDJK2-K14-GFPNLS, and AAVDJK2-K16-GFPNLS plasmids were collaterally transfected into 293 AAV cells in eight flasks. Cells were harvested and processed to PEG-concentrated aliquots (100 µl for each flask), and the titer was measured by qPCR [2][16].

### Image analyses

Imaging was performed using a Stereoscope Zeiss AXIO Zoom.V16, Olympus VS-200 Slide Scanner, and Olympus IX73 Inverted LED Fluorescence Microscope. Analyses were carried out using ImageJ software version 1.53.

## Results

### Generation and evaluation of the gene transduction efficiency of AAVDJ variants in cultured KCs

Site-directed mutagenesis of AAVDJ was employed to generate four AAVDJ variants with mutations reported to increase tropism of AAV2 for KCs [10]. The gene transduction efficiencies of these AAVDJ variants in cultured mouse and human KCs were compared with those of seven AAVs reported previously as optimal vectors for gene transduction of KCs (AAV2 and four of its variants [10], AAVDJ [11], and AAV6 [19]) (Fig. 1a).

**Fig. 1.**
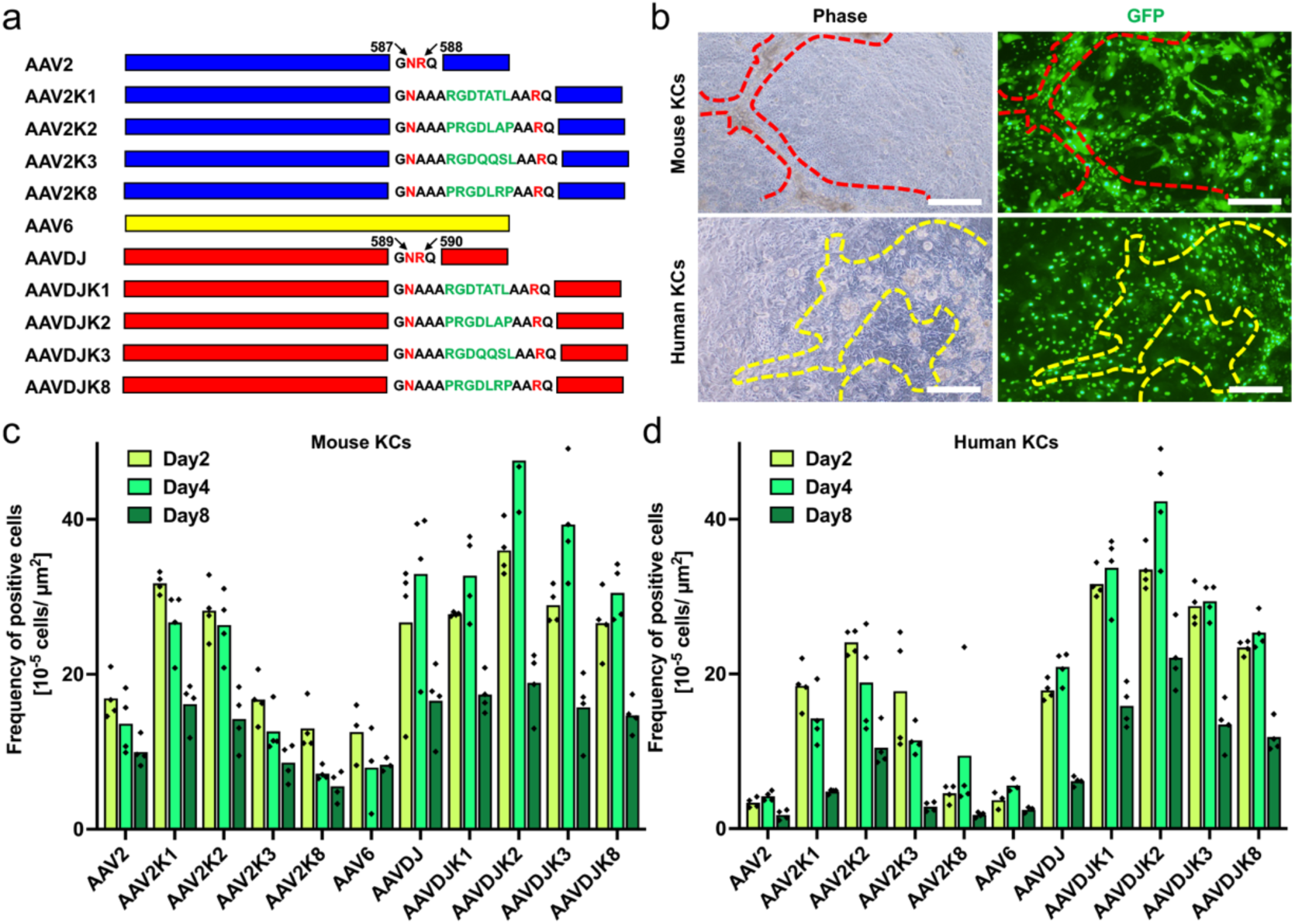
AAVDJK2 is effective in cultured mouse and human keratinocytes. a) Schematic of the amino acid sequence of modified AAVDJ and AAV2 with mutations and AAV6 prepared for comparison. Additional peptide was inserted just before Arg588 in AAV2 capsid and Arg590 in AAVDJ capsid. b) Appearance of AAVDJK2-treated mouse and human keratinocytes on feeders. Dotted red line indicates the colony border in mouse KC culture. Dotted yellow line indicates the colony border in human KC culture. Scale bar = 100 µm. c) GFPNLS-positive cell counts of AAV-treated mouse KCs. Mean and distribution of the data are shown.d) GFPNLS-positive cell numbers of AAV-treated human KCs. Mean and distribution of the data are shown.

AAVs encoding green fluorescent protein with a nuclear localization signal (GFPNLS) were used to assess gene transduction efficiency because nucleus-localized fluorescence facilitates distinguishing gene expression from background signals [7]. KCs cultured on feeder cells form colonies with a defined edge [2][20]. At 2, 4, and 8 days after addition of AAVs, the number of GFPNLS-positive cells in the KC colony area was counted (Fig. 1b). For both mouse and human KCs, gene transduction efficiency, as assessed by the number of GFPNLS-positive cells per KC colony area, was consistently highest with a type of mutant AAVDJ termed AAVDJK2 (Fig. 1c, 1d). AAVDJK2 demonstrated approximately twice the gene transfer efficiency of previously known capsids on keratinocytes *in vitro*.

## Evaluation of gene transduction efficiency of AAVDJK2 in cultured mesenchymal cells

To evaluate the specificity of AAVDJK2 for skin, the gene transduction efficiency in skin and soft tissue-derived mesenchymal cells, mouse adipose-derived stromal cells (mASCs), and human dermal fibroblasts (hDFs), was examined in comparison with AAV2, AAV2K2, and AAVDJ. We found that the gene transduction efficiency of AAV2 differed between mASCs and hDFs, the highest tropism towards both mouse and human mesenchymal cells specific to the original AAVDJ [2] was lost in AAVDJK2 (Fig. 2a, 2b), which increased specificity of AAVDJK2 for epidermal cells.

**Fig. 2.**
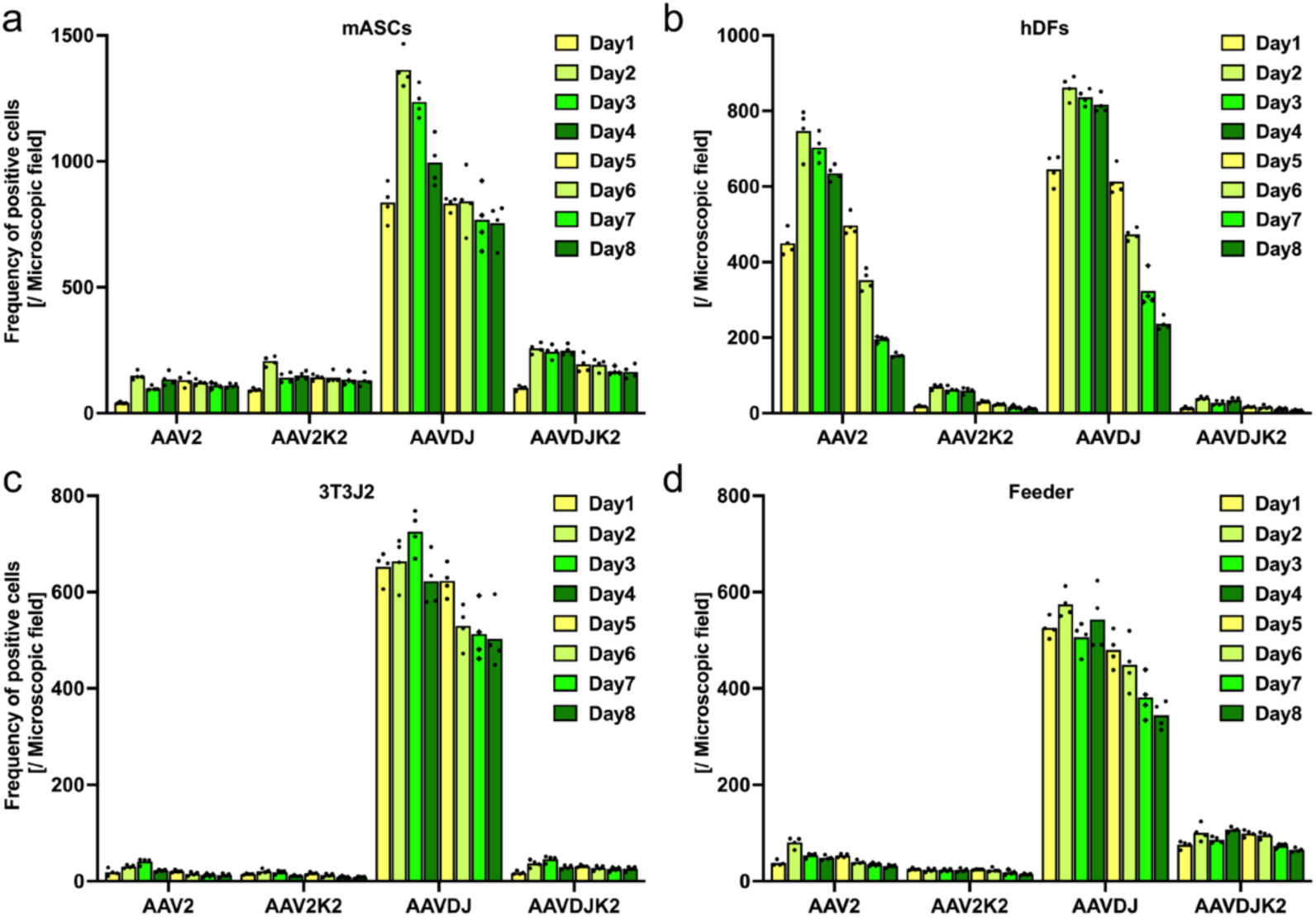
AAVDJK2 reduces AAVDJ-specific tropism for mesenchymal cells. a) GFPNLS-positive cell numbers of AAV-treated mouse adipose-derived stromal cells. Mean and distribution of the data are shown. b) GFPNLS-positive cell numbers of AAV-treated human dermal fibroblasts. Mean and distribution of the data are shown. 3) GFPNLS-positive cell numbers of AAV-treated 3T3J2 cells. Mean and distribution of the data are shown. 4) GFPNLS-positive cell number of AAV-treated feeder cells (mitomycin treated 3T3J2 cells). Mean and distribution of the data are shown

For future *in vitro* research and development, we also examined the efficiency of gene transfer by the 4 AAVs on mitomycin untreated and treated 3T3-J2 cells, a mesenchymal cell line derived from fetal mice and best known as a feeder for keratinocytes. A trend similar to that observed in mASC, with only AAVDJ showing high gene transfer efficiency, was also observed in 3T3-J2 cells (Fig. 2c, 2d).

### Evaluation of the nanoparticle distribution after intradermal injection

To evaluate efficiency and specificity *in vivo*, intradermal injection was used to deliver AAVs as a simple, consistent, reproducible, and highly accessible method. Although intradermal injection is a very common methodology, to our knowledge, no studies have examined the *in vivo* distribution of an injected drug after injection into mouse skin. Therefore, after injection, we investigated the distribution of fluorescent silica nanoparticles with a diameter of 30 nm, which was similar to that of AAVs [7][21]. The distribution of fluorescent silica nanoparticles was examined histologically at 0, 1, and 3 hours after intradermal injection (Fig. 3a–c). We found that only a small fraction of the nanoparticles diffused into the epidermis over time (Fig. 3d). This result suggested that only a small portion of administered AAVs entered the epidermis.

**Fig. 3.**
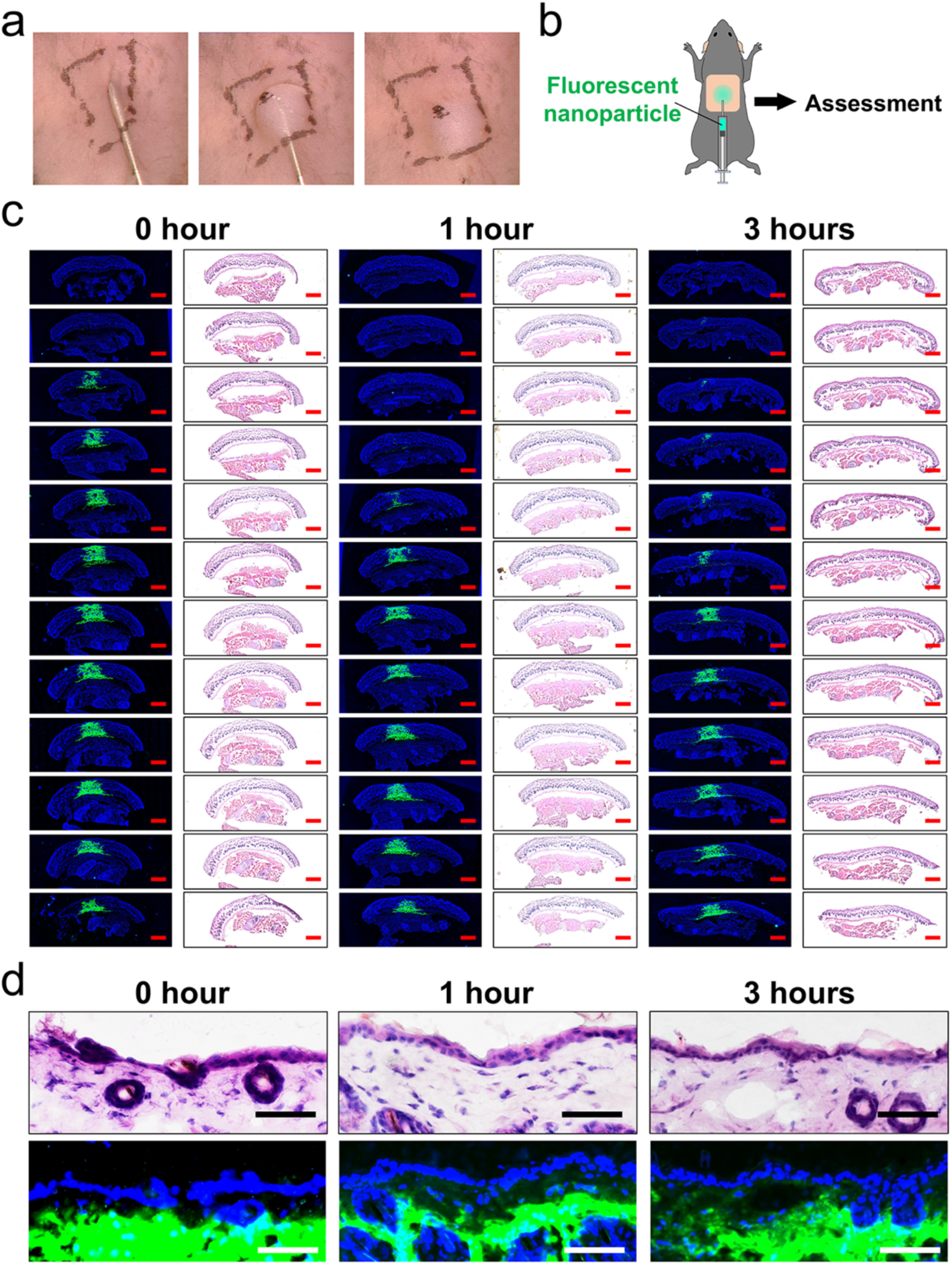
Most injected nanoparticles diffuse into the dermis and subdermal tissues after intradermal injection. a) Schematic of fluorescent nanoparticle injection. b) Appearance of the intradermal injection. Black dotted line was designed as a 1 cm square rectangle. c) Representative findings of three-dimensional histological analysis of serial sections of skin samples. Epidermal diffusion was evaluated in the section with the broadest nanoparticle diffusion. Red scale bar = 1 mm. d) Representative magnified results of histological analysis of skin samples. Over time, diffusion of the injected nanoparticle towards the epidermis was low. Scale bar = 50 µm.

### Evaluation of AAVDJK2 *in vivo*

Assuming the inefficient properties of intradermal injection into mouse skin, the gene transduction efficiency of AAVDJK2 after intradermal injection into mouse skin was evaluated in comparison with AAVDJ. The time of sample collection was 2 days after injection because, in preparatory *in vivo* assessments, the frequency of fluorescence-positive cells was higher than at 4 and 7 days after injection, which is consistent with the rapid turnover cycle (8–10 days) of the mouse epidermis [22]. To minimize variances caused by the administration, a mixture of 1×10^11^ GC (gene copies) of AAVDJK2-CAG-GFPNLS and 1×10^11^ GC AAVDJ-CAG-mCherryNLS was prepared as a 35 µl solution and injected intradermally into the back skin of mice (Fig. 4a) (n=6). After confirming the position of the center of AAV delivery by fluorescence stereoscopic observation (Fig. 4b), specimens were collected. Frozen serial sections were prepared at 200 µm intervals, nuclei were stained with DAPI, and samples were imaged using the slide scanner.

**Fig. 4.**
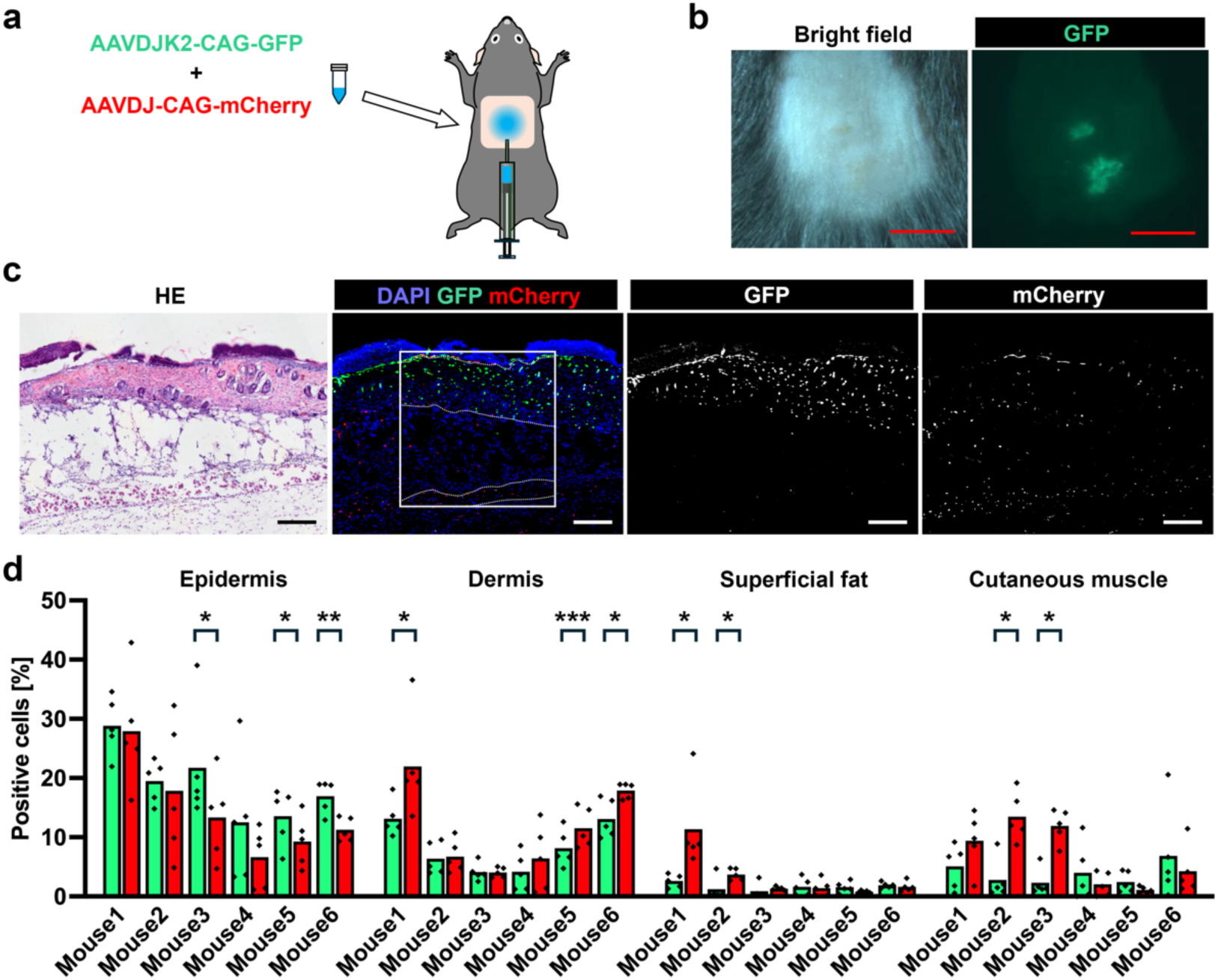
AAVDJK2 has high epidermal specificity in mouse skin. a) Schematic of intradermal injection of mixtures of AAVDJK2-CAG-GFPNLS and AAVJD-CAG-mCherryNLS. b) Confirmation of the position of AAV delivery by fluorescence stereoscopy. Red scale bar = 5 mm. c) Representative hematoxylin and eosin-stained sections and DAPI-stained fluorescence sections. The square indicates the areas used for deep learning-based image analysis. Dotted line indicates the boundaries of skin layers used for layer-specific analyses (epidermis, dermis, superficial fat, cutaneous muscle, and deep fat). Scale bar = 200 µm. d) Percentage of transgene expression-positive cells in each layer in six animals. Differences between groups were analyzed by the paired Student’s t-test. *P<0.05, **P<0.01, and ***P<0.001 (n=6).

Nucleus-localized fluorescence was detected from the superficial to deep layer around the injection site (Fig. 4c). The expression efficiency of AAVDJK2-CAG-GFPNLS and AAVDJ-CAG-mCherryNLS in each skin layer (epidermis, dermis, superficial fat, and cutaneous muscle) was quantified by automated and unbiased analysis.

Considerable intersectional variation and fluorescence detection were observed, but significant differences in the expression patterns of GFPNLS between AAVDJK2 and AAVDJ were found in several animals per layer. The number of AAVDJK2-delivered, GFPNLS-positive cells was higher than that of AAVDJ-delivered, mCherryNLS-positive cells in the epidermis, and the opposite trend was observed in deeper layers (Fig. 4d). Thus, the higher epidermal specificity of AAVDJK2 suggested by *in vitro* analyses was confirmed in mouse skin *in vivo*.

### Development and evaluation of tissue-specific promoters

Another approach to increase the efficiency and specificity of gene delivery to specific cells/tissues is the use of tissue-specific promoters.

With reference to the reported *KRT14* (K14) promoter [13], enhancer, and minimal promoter elements [14], we generated constructs containing GFPNLS under the control of the human K14 promoter, K14 short promoter with endogenous minimal promoter, and original hybrid short promoter composed of the K14 enhancer and engineered potent core promoter, SCP3 [15]. For each construct, another version with multiple single nucleotide polymorphisms in the promoter sequence for the *KRT16P5* pseudogene used in the commercially available keratinocyte differentiation reporter vector was also prepared (Fig. 5a). Examination of the GFPNLS-positive rate of mouse KCs and mASCs treated with AAVDJK2-packaged AAVs with each promoter and the ubiquitous CAG promoter revealed that the K14 SCP3 promoter had the best gene transduction efficiency in keratinocytes, whereas expression in mASCs was maintained at relatively low levels (Fig. 5b–e). The *In vivo* gene transduction efficiency of AAVDJK2-CAG-GAPNLS and AAVDJK2-K14 SCP3-GFPNLS was tested (n=3, each) and it was confirmed that the use of the K14 SCP3 promoter increased the specificity for the epidermis without apparent loss of the gene expression efficiency of the CAG promoter (Fig. 6).

**Fig. 5.**
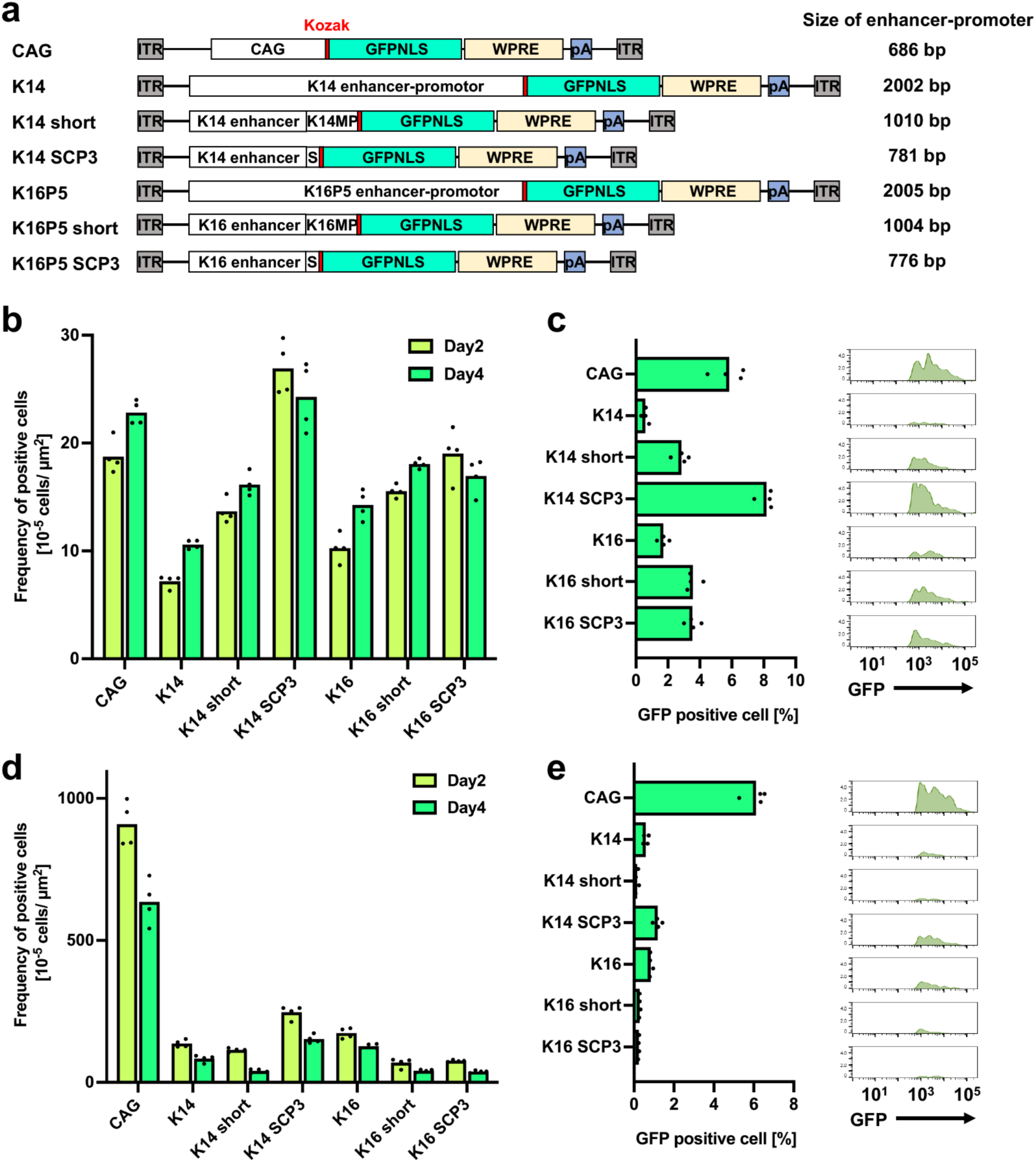
K14 SCP3 promoter enhances keratinocyte specificity *in vitro*. a) Schematic of the tissue-specific promoter vector. K14MP, *KRT14* minimal promoter. K16MP, *KRT16 pseudogene 5* minimal promoter. S, SCP3 core promoter. pA, polyA signal. WPRE, Woodchuk hepatitis virus Posttranscriptional Regulatory Element. b) GFPNLS-positive cell counts of AAV-treated mouse keratinocytes. Mean and distribution of the data are shown. c) Flow cytometric analysis of AAV-treated mouse keratinocytes. Mean and distribution of the data and representative histograms are shown. d) GFPNLS-positive cell counts of AAV-treated mouse adipose-derived stromal cells. Mean and distribution of the data and representative histograms are shown. e) Flow cytometric analysis of AAV-treated mouse adipose-derived stromal cells. Mean and distribution of the data and representative histograms are shown.

**Fig. 6.**
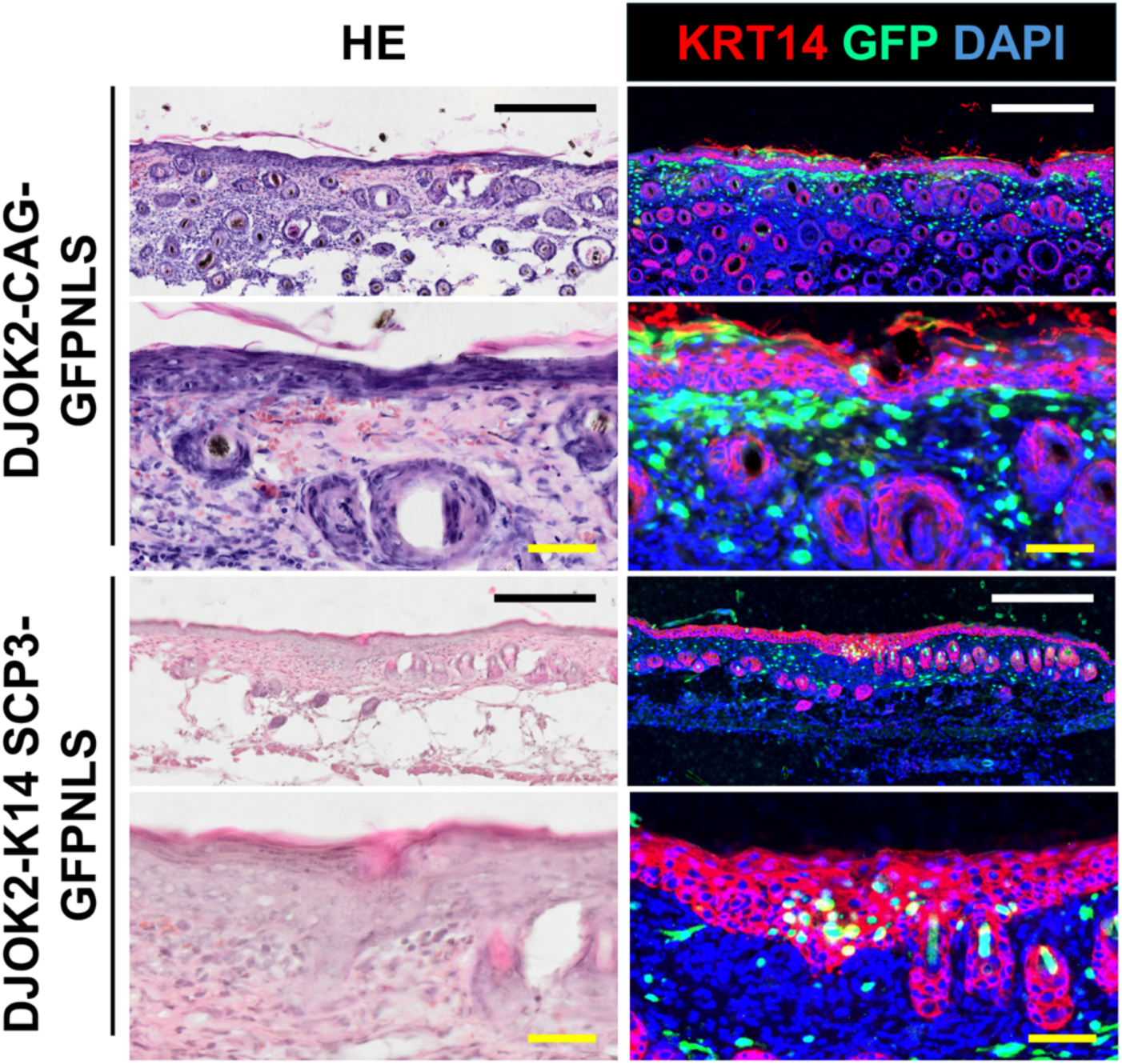
K14 SCP3 promoter enhances epidermal specificity *in vivo*. Representative hematoxylin and eosin-stained sections and immunofluorescence sections of mouse skin after intradermal injection of AAVDJK2-CAG-GFPNLS and AAVDJK2-K14 SCP3-GFPNLS. Black and white scale bar = 300 µm. Yellow scale bar = 100 µm.

### Effects of the capsid and promoter on AAV production

In our experiences with repeated production of AAVs with various capsids, promoters, and genes [2][7][23], AAV production using the same procedure is substantially affected by the type of capsid, promoter, and gene. We examined AAV production of various types of capsids and promoters, and confirmed significant differences (Supplementary Fig.1), suggesting that AAV production is a practical reason for selecting AAV types, especially when used for experiments.

## Discussion

The newly developed AAVDJ variant capsid termed AAVDJK2 showed higher gene transduction efficiency in mouse and human KCs *in vitro* than other AAVs reported to be appropriate for KCs. The use of high doses results in stronger phenotypes. However, most AAV vectors cause non-specific effects [24]. In this context, the specificity of gene expression at target tissues, minimizing off-target effects, is important for *in vivo* applications [7]. Changing the tropism of an AAV to a specific cell type may affect its interaction with other cell types. Therefore, we investigated the effect of the mutagenesis on the efficiency of gene transduction in representative skin-derived non-epidermal cells. As expected, the mutagenesis reduced the tropism for both mASCs and hDFs, implying acquisition of epidermal specificity in skin.

To evaluate the utility of AAVDJK2 for *in vivo* gene transduction of the epidermis, we conducted experiments using mouse skin by intradermal injection, followed by analysis of the distribution of injected particles. Intradermal injection was not an efficient administration method for the mouse epidermis. However, in the central region, AAVDJK2 consistently achieved a positive rate of approximately 20% of cells in all epidermal layers. The comparison between AAVDJK2 and AAVDJ was not as clear as that in the *in vitro* experiment, but as suggested, the higher gene transfer efficiency and specificity of AAVDJK2 were confirmed. In current capsid engineering, peptide modifications were targeted at the heparin-binding region, situated on the outermost surface of AAV2 and AAVDJ capsids in three dimensions, exerting a significant influence on cellular targeting by strongly binding to heparin sulfate proteoglycans on cell surfaces [10] [25]. As demonstrated for AAV2 [10], these modifications in AAVDJ may have resulted in stronger binding of the capsid to αvβ8 integrin on the surface of keratinocytes, thereby increasing infectivity. Thus, the differential expression of sites across different capsids is not attributed to changes in AAV distribution but rather to variations in the types of cells that AAV can effectively target, as confirmed by *in vitro* assessments.

The thickness of the mouse dermis is much thinner (100–200 µm) than that of the human dermis (several millimeters) [26]. Moreover, the subcutaneous fat layer under the mouse dermis is very rough and flexible. Therefore, injected drugs disperse easily subcutaneously, making it difficult for drugs injected into the dermis to achieve effective delivery to the epidermis. Conversely, intradermal injection into human skin can be performed more consistently. Furthermore, various types of specialized methods, such as microneedles, fractional laser, and other methods that use physically created small holes and chemical drug carriers to enhance penetration, are being developed for epidermis-specific drug delivery in humans [27] [28] [29] [30]. Human skin is likely to be more amenable than mouse skin in terms of drug delivery to the epidermis, which should be considered when evaluating results from mouse skin as a model.

In this study, to achieve high specificity using AAVDJK2, we developed K14 SCP3, which combined a *KRT14*-derived enhancer with a recently reported short core promoter, and confirmed *in vitro* that it had higher gene expression and specificity in KCs than CAG (ubiquitous strong promoter) and previously reported specific promoters. This enhancer-promoter sequence is relatively short (781 bp) and is expected to be effective in AAVs because of their limited capacity (approximately 4.7 kb).

The amount of AAV production varies with the capsid and promoter, even when the same gene is mounted and the same procedure is used for production. This is especially important to note for experimental use because the efficiency of AAV production is very important in terms of cost and labor.

In the current study, we did not analyze gene transfer efficiency and gene expression intensity for specific terminal differentiation states of the epidermis because the accuracy of the fluorescent signal detection decreases as the terminal differentiation progresses. The epidermis is a tissue with a strong background signal, especially in the superficial layers and near the stratum corneum. The nuclear localization signal becomes unreliable due to denucleation during terminal differentiation. We anticipate that in most applications targeting keratinocytes and epidermis, progenitor cell fractions will be targeted. However, for special applications, more focused studies, such as calcium-added differentiation induction in non-serum culture, are envisioned.

Many skin disorders are caused by single gene mutations in keratinocytes. Some types of skin diseases, such as epidermolysis bullosa, are clinically treated by ex vivo gene therapies [31][32]. Recently, other types of *in vivo* topical approaches are emerging [33] [34]. Although AAVs have not received much attention as an *in vivo* topical gene transfer method for the epidermis, AAVs have some unique advantages, such as capsid tropism. Along with ongoing development of various epidermal specific delivery systems [27] [28] [29] [30], our optimized AAV system for the epidermis/ keratinocytes is useful for developing new epidermis-targeted gene therapies.

## Acknowledgments

We thank Mitchell Arico from Edanz (https://jp.edanz.com/ac) for editing a draft of this manuscript.

## Abbreviations

KC: keratinocyte
AAV: adeno-associated virus
PBS: phosphate-buffered saline
EDTA: ethylenediaminetetraacetic acid
PFA: paraformaldehyde
DAPI: 4ʹ,6-diamidino-2-phenylindole
GFPNLS: green fluorescent protein with a nuclear localization signal
mCherryNLS: mCherry with a nuclear localization signal
mASCs: mouse adipose-derived stromal cells
hDFs: human dermal fibroblasts
K14: *KRT14*
K16P5: *KRT16* pseudogene 5

---

**Supplementary Fig. 1.**
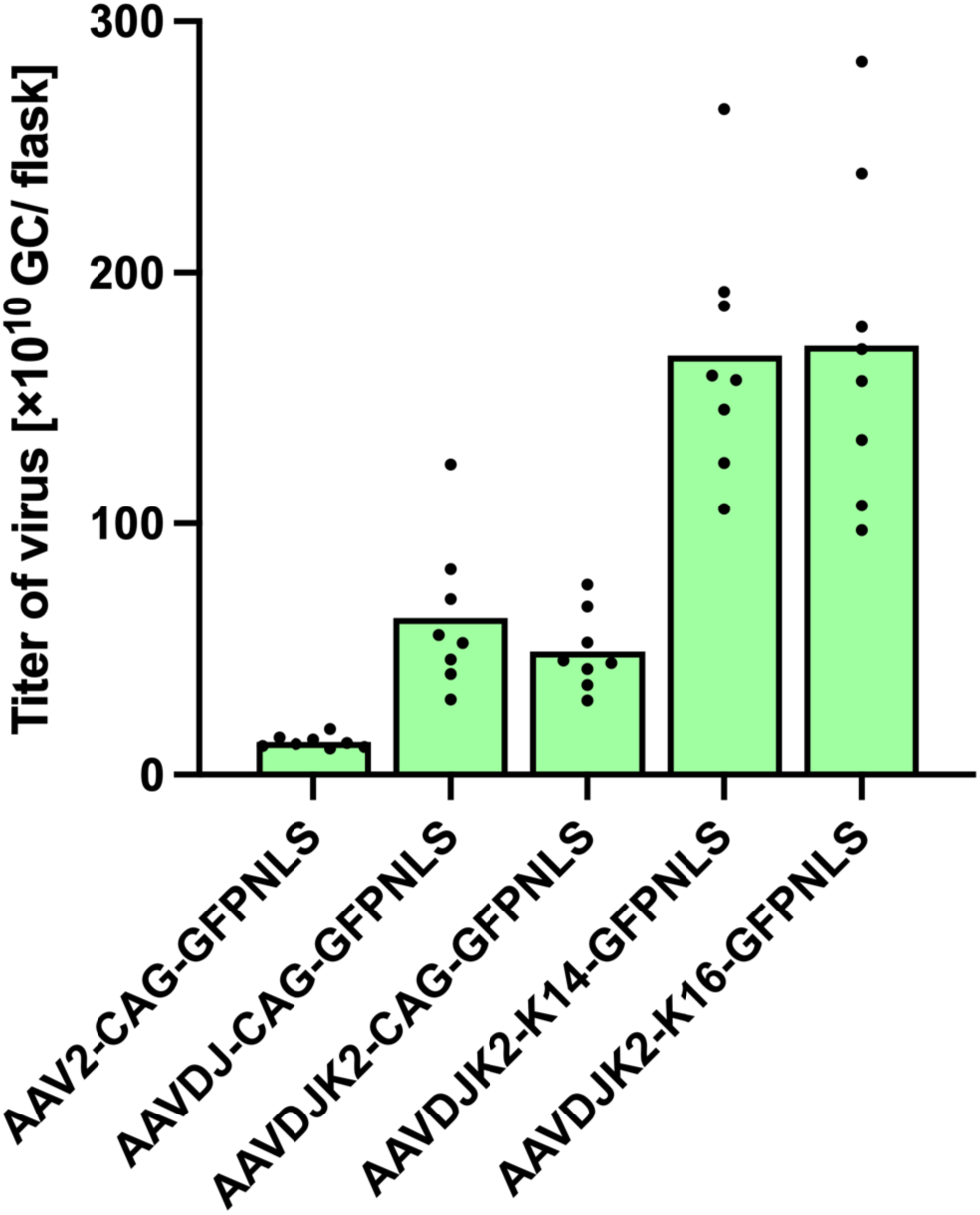
AAV yields are influenced by the capsid and promoter. AAV yields of GFPNLS-expressing AAVs with various capsids and promoters were evaluated. AAVDJ and AAVDJK2 were superior to AAVs in terms of packaging CAG-GFPNLS AAVs, whereas K14 and K16 promoters were superior to GFPNLS-expressing AAVDJK2 AAVs.

